# Novel RT-ddPCR Assays for determining the transcriptional profile of SARS-CoV-2

**DOI:** 10.1101/2021.01.12.425991

**Authors:** Sushama Telwatte, Nitasha Kumar, Albert Vallejo-Gracia, G. Renuka Kumar, Chuanyi M. Lu, Melanie Ott, Joseph K. Wong, Steven A. Yukl

**Author notes:** Corresponding author: Steven Yukl; San Francisco VA Medical Center, 4150 Clement St, 111W3, San Francisco, CA 94121, USA.

## Abstract

The exact mechanism of coronavirus replication and transcription is not fully understood; however, a hallmark of coronavirus transcription is the generation of negative-sense RNA intermediates that serve as the templates for the synthesis of positive-sense genomic RNA (gRNA) and an array of subgenomic mRNAs (sgRNAs) encompassing sequences arising from discontinuous transcription.

Existing PCR-based diagnostic assays for SAR-CoV-2 are qualitative or semi-quantitative and do not provide the resolution needed to assess the complex transcription dynamics of SARS-CoV-2 over the course of infection. We developed and validated a novel panel of specially designed SARS-CoV-2 ddPCR-based assays to map the viral transcription profile. Application of these assays to clinically relevant samples will enhance our understanding of SARS-CoV-2 replication and transcription and may also inform the development of improved diagnostic tools and therapeutics.

**Highlights:**

- We developed a novel panel of 7 quantitative RT-ddPCRs assays for SARS-Cov-2
- Our panel targets nongenic and genic regions in genomic and subgenomic RNAs
- All assays detect 1-10 copies and are linear over 3-4 orders of magnitude
- All assays correlated with the clinical Abbott SARS-CoV-2 Viral Load Assay
- Clinical samples showed higher copy numbers for targets at the 3’ end of the genome

## 1. Introduction

The etiologic agent responsible for the ongoing COVID-19 pandemic, identified as Severe Acute Respiratory Syndrome Coronavirus 2 (SARS-CoV-2),^1,2^ is an enveloped virus with a positive-sense, single-stranded RNA genome of ~30 kb and is a member of the β-coronavirus genus. SARS-CoV-2, which is the seventh coronavirus known to infect humans, shares approximately 50% sequence homology with MERS and 79% sequence homology with SARS-CoV^3^ but appears to be more closely related to the SARS-like bat coronaviruses RmYN02 from R. malayanus and RaTG13 from R. affinis (93.3% and 96.1% sequence identity, respectively),^4^ though its origin is, to date, unsettled^5,6^.

The exact mechanism of SARS-CoV-2 replication and transcription is not fully understood; however, a hallmark of coronavirus transcription and other viruses of the order *Nidovirales* is the generation of negative-sense RNA intermediates that serve as the templates for the synthesis of positive-sense genomic RNA (gRNA) and an array of subgenomic RNAs (sgRNAs), which arise from discontinuous transcription and encompass sequences from both ends of the genome ^7,8^ (Fig. 1). Following cell entry, SARS-CoV-2 genomic RNA is transcribed and translated to generate the nonstructural proteins (NSP) from the two open reading frames (ORF), ORF1a and ORF1b^8^, a process thought to involve the virus replication complex, transcription-regulating sequences (TRSs), the N protein, and double membrane vesicles in the cytoplasm of infected cells^9–11^. During the synthesis of the negative strand RNA, sgRNAs arise from a template switch that adds a copy of the ‘leader’ sequence (~70 nucleotides in the 5’ untranslated region [UTR] containing a short transcription-regulating sequence [TRS] at the 3’ end) to the ‘body’ sequence derived from one of various genes in the 3’ third of the genome (including genes for structural proteins)^12–14^. Transcription of the sgRNAs is likely regulated by TRS sequences in the leader sequence and upstream of 3’ genes^9^, and may allow for greater expression of certain viral genes.

**Figure 1.**
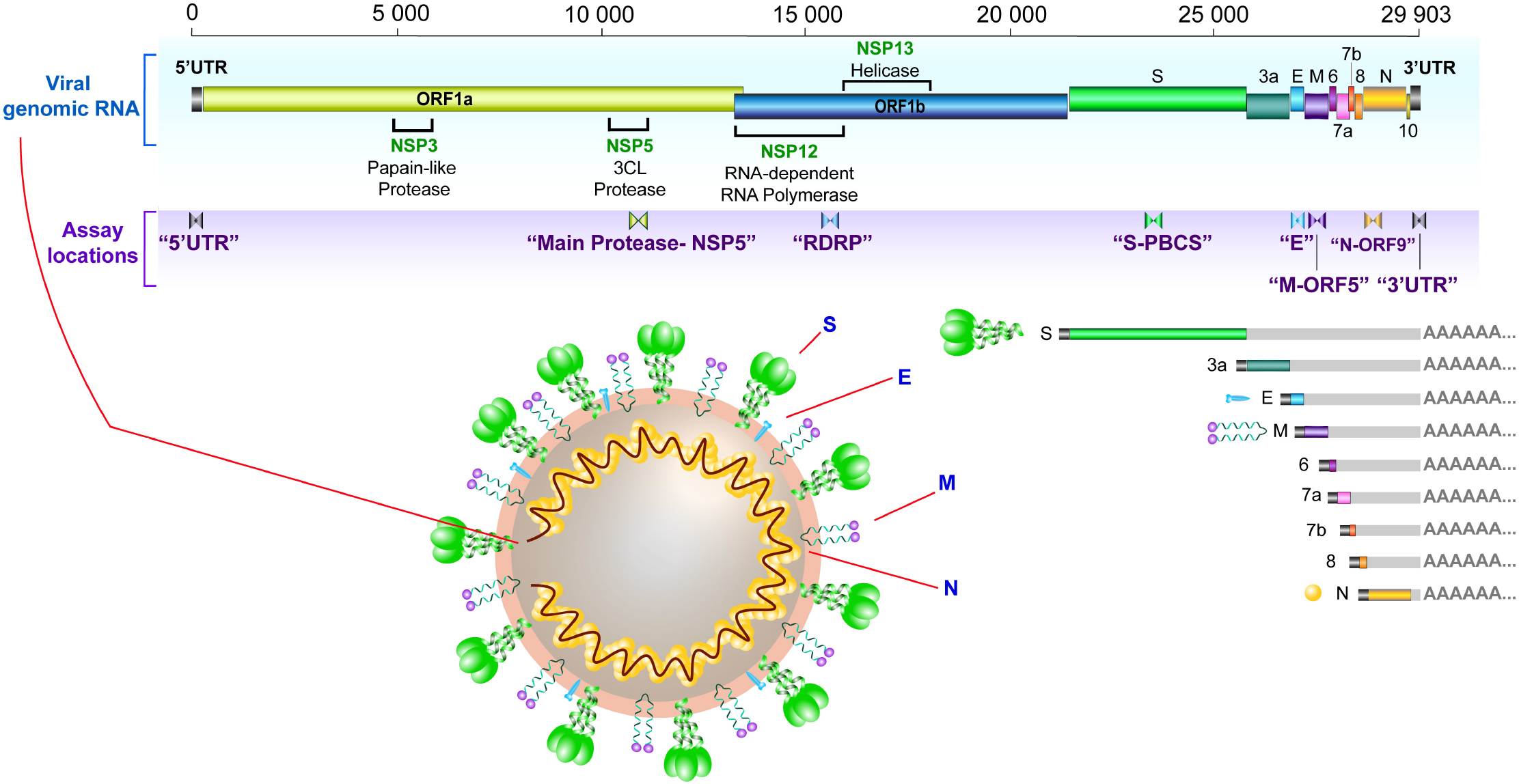
Schematic presentation of SARS-CoV2 genome organization, virion structure and canonical sgRNAs. SARS-CoV-2 encodes two large genes, ORF1a (yellow) and ORF1b (blue), which encode 16 non-structural proteins (NSP1–NSP16). The structural genes encode the structural proteins, spike (S; green), envelope (E; blue), membrane (M; purple), and nucleocapsid (N; gold). Assay locations of each assay designed for this study are indicated. Virion structure and canonical subgenomic RNAs produced by SARS-CoV-2 are shown in the lower panel (S, 3a, E, M, 6, 7a, 7b, 8 and N).

A recently published study confirms that a similar mechanism exists for SARS-CoV-2 to generate nine canonical sgRNAs distinct from genomic RNA^8^ (Fig. 1). For other coronaviruses, sgRNAs encode virulence factors such as proteins that directly cause lesions^15^ or indirectly inhibit immune responses^16^. Incorporation of 5’UTR sequences into the capped subgenomic mRNA templates of SARS-CoV may confer resistance to cleavage by viral nsp1 protein^17^, which typically inhibits host gene expression by degradation of host mRNA^18–20^. For positive-sense RNA viruses, sgRNAs act as messengers for expression of structural proteins or proteins related to pathogenesis and can regulate the transition between translation and virion production^21^. The various roles of sgRNAs in SARS-CoV-2 infection and pathogenesis remain to be elucidated, but the rapid accumulation and persistence of sgRNAs following infection may also contribute to disease progression.

Understanding the viral dynamics of SARS-CoV-2 and the host response are essential in devising strategies to develop antiviral treatments or vaccines and curb new infections. Existing PCR-based diagnostic assays for SAR-CoV-2, which are interpreted in a qualitative or semi-quantitative manner (positive, negative or indeterminate) and target only 1-2 viral regions, do not distinguish between genomic and subgenomic RNA or account for possible differences between the RNA copy numbers of various viral genes, which may depend on the degree to which they are transcribed as various sgRNAs and the degree to which the sample includes virion or cell-associated RNA. Molecular assays that can quantify different viral genes found in genomic and sgRNA species will have utility in charting the extent of viral replication and changes in SARS-CoV-2 transcription over the course of infection.

We have devised a novel panel of seven ddPCR-based assays that target various conserved regions of SARS-CoV-2 RNA, including the 5’ and 3’ untranslated regions, non-structural genes that are only found in full length (genomic) RNA and structural genes that are also contained in different sgRNAs (Fig.1 and Table 1).

**Table 1.** SARS-Cov2 ddPCR assay panel for assessing patient samples

| Assay Name | RNA Target | Detects |
| --- | --- | --- |
| **5'UTR** | 5' untranslated region | Genomic RNA |
| **Main Proteinase-NSP5** | Main Proteinase | Genomic RNA |
| **RdRp** | RNA-dependent RNA Polymerase | Genomic RNA |
| **S-PBCS** | <u>P</u>oly<u>b</u>asic <u>c</u>leavage <u>s</u>ite of the surface (S) glycoprotein | Genomic/subgenomic |
| **M-ORF5** | Membrane glycoprotein | Genomic/subgenomic |
| **N-ORF9** | Nucleocapsid | Genomic/subgenomic |
| **3'UTR** | 3' untranslated region | Genomic/subgenomic |

We selected genes encoding two non-structural proteins [Main Proteinase (NSP5) and RNA dependent RNA polymerase (RdRp-NSP12)] and four major structural proteins [Spike glycoprotein (S), envelope (E), membrane (M), and nucleocapsid (N)] that are known to serve critical functions in SARS-CoV-2 infection. For the spike protein, in which notable mutations have emerged^22^, we designed a primer/probe set to target the short, highly-conserved ‘polybasic cleavage site’ (‘S-PBCS’) of SARS-CoV-2 which is functionally cleaved to yield the S1 and S2 subunits^23^, in a similar manner to the hemagglutinin (HA) protein of avian influenza viruses (AIVs)^24^. In AIVs, the insertion or substitution of basic amino acids at the HA cleavage site is associated with enhanced pathogenicity^25,26^.The SARS-CoV-2 PBCS allows effective cleavage by host furin and other proteases^5^, and may potentially enhance its infectivity in humans and distinguish it from related animal coronaviruses^4,5,27^. Elucidating the granular detail of SARS-CoV-2 transcription could help us to understand how the virus replicates and how it may evade human immune defenses. Detailed mapping of the expressed viral transcripts across times and cell types is essential for further studies of viral gene expression, mechanisms of replication, and probing host-viral interactions involved in pathogenicity.

## 2. Materials and methods

### 2.1 Primer design and selection

Multiple primer and probe sets were designed to target various regions of SAR-CoV-2, including untranslated regions that likely play an important role in regulating transcription (5’ and 3’ untranslated regions [UTR]), non-structural genes found only in genomic RNA (main protease [NSP5; ORF1a], RNA-dependent RNA polymerase [RdRp; ORF1b]), and structural genes that may also be found in various sgRNAs (spike [S] protein [ORF2] polybasic cleavage site [PBCS], membrane [M] glycoprotein [ORF5], and nucleocapsid [N] protein [ORF9]). Primers/probes were designed using the Primer Quest® Tool (Integrated DNA Technologies, Coralville, IA). A multiple sequence alignment was performed using Clustal Omega^28^, encompassing complete sequences of 86 SARS-CoV-2 isolates from all geographical locations and all sequences available from the US on 3/14/2020. Reference sequences of other coronaviruses, including SARS-CoV (NC_004718.3), MERS-CoV (NC_019843.3), HCoV-229E (NC_002645.1), HcoV-NL63 (NC_005831.2), HcoV-OC43 (NC_006213.1), and HcoV-HKU1 (NC_006577.2), were included in the alignment to exclude primer sets with significant overlap with non-SARS-CoV-2 sequences. Two primer/probe sets that aligned to all SARS-CoV-2 isolates but had 1 or more mismatch with SARS-CoV and greater than 5 mismatches with MERS-CoV, HCoV-229E, HCoV-NL63, HCoV-OC43, HCoV-HKU1 were selected for each region (Table 1). A sequence similarity analysis using Basic Local Alignment Search Tool (BLAST)^29^ found no significant similarity in any primer or probe to human sequences.

### 2.2 Validations using plasmid DNA

Plasmid constructs containing the regions of interest (5’UTR, 3’UTR, Main Proteinase, M gene, N gene, S protein, and a 528nt fragment of RdRp) were designed in pBluescript KS(+) (Bio Basic Inc., Ontario, Canada) to enable assay validations using DNA and for use in *in vitro* transcription reactions to generate viral RNA for standards. Plasmid concentrations were quantified using ultraviolet (UV) spectrophotometry (NanoDrop ND-1000 instrument, Thermo Fisher) and the molecular weights were used to calculate the number of molecules per μL. Extracted PBMC from a healthy donor (150-200 ng/well) and H_2_O were included as negative controls for each assay.

Each primer and probe set was tested using droplet digital PCR (ddPCR), as performed using the QX100 system (Bio-Rad). Droplet digital PCR was chosen because it enables “absolute” quantification, it is relatively less dependent on PCR efficiency (which may be reduced by sequence mismatches or inhibitors), and it may be more precise than quantitative PCR (qPCR) at low copy numbers^30^. Plasmid DNA was added to ddPCR wells at expected inputs of 1-10^3^ copies/well in duplicate (1000 and 100 copies) or quadruplicate (10 and 1 copy). Each reaction consisted of 20 μL per well containing 10 μL of ddPCR Probe Supermix (no deoxyuridine triphosphate), 900 nM of primers, 250 nM of probe, and 5 μL of plasmid DNA. Droplets were amplified using a Mastercycler® nexus (Eppendorf, Hamburg, Germany) with the following cycling conditions: 10 min at 95°C, 45 cycles of 30 s at 95°C and 59°C for 60 s, and a final droplet cure step of 10 min at 98°C. Droplets were read and analyzed using the QuantaSoft software in the absolute quantification mode.

### 2.3 Validations using synthetic RNA

*In vitro* transcribed (IVT) RNA standards were generated from the aforementioned plasmids using the T7 RiboMAX™ Express Large-Scale RNA Production System (Promega, Madison, WI). The concentration of each IVT RNA standard was measured by Nanodrop and the molecular weight was used to calculate the expected number of molecules per μL. The length, integrity, and concentration of each IVT standard were confirmed using the Agilent Bioanalyzer RNA 6000 Nano assay (Agilent, Santa Clara, CA) prior to dilution in nuclease-free water to working concentrations.

A reverse transcription (RT) reaction was performed in 50 μL containing 5 μL of 10× SuperScript III buffer (Invitrogen), 5 μL of 50 mM MgCl2, 2.5 μL of random hexamers (50 ng/μL; Invitrogen), 2.5 μL of 50 μM poly-dT15, 2.5 μL of 10 mM deoxynucleoside triphosphates (dNTPs), 1.25 μL of RNAseOUT (40 U/μL; Invitrogen), and 2.5 μL of SuperScript III RT (200 U/μL; Invitrogen). Although the IVT standards were not polyadenylated, reverse transcription was performed with both random hexamers and poly-dT because we anticipated that these assays would be applied to clinical samples containing long polyadenylated SARS-CoV-2 RNAs, for which the combination of poly-dT plus random hexamers may reduce bias towards reverse transcription of any one region (as can be seen with specific reverse primers), the 5’ end (as would be expected with random hexamers), or the 3’ end (as would be expected with poly-dT).

IVT RNA standards were added to RT reactions at inputs of 1, 10, 10^2^, 10^3^, and 10^4^ copies per 5 μL (2 replicate RT reactions for each input). RT reactions were performed in a conventional thermocycler at 25.0°C for 10 min, 50.0°C for 50 min, followed by an inactivation step at 85.0°C for 5 min. Undiluted RT product (5 μL) was added to ddPCR reactions (total volume of 20 μL) and ddPCR was performed as described for ‘Validations using plasmid DNA’. Primer-probe sets for each target region were tested head-to-head using this approach. Based on performance of each primer-probe set using plasmid DNA and IVT RNA, one primer/probe set for each region was selected for further testing.

To determine the robustness of our approach, in addition to testing each assay with varying RNA copy inputs (each with two replicate RT reactions per input and replicate ddPCR wells for each RT), we performed repeat, independent experiments using the same parameters to confirm each assay’s efficiency and sensitivity (n=4 for N-ORF9, CDC_N1, and CDC_N2; n=3 for 5’UTR, 3’UTR; and n= 2 for all others). No data were excluded as outliers.

### 2.4 Validations using SARS-CoV-2 virion RNA

Vero CCL-81 kidney epithelial cells, derived from *Cercopithecus aethiops*, were infected with SARS-CoV-2 (Isolate: USA-WA1/2020) at an MOI of 0.003 (250 000 cells/well). Cells were incubated for 72 hours at 37°C/5% CO_2_ and harvested. Viral supernatant was clarified by 2 centrifugation steps (180 *xg*, 5 min) and added directly to 1mL TRI reagent (Molecular Research Center Inc.). Total RNA was extracted using TRI reagent, including the addition of polyacryl carrier (2.5μL). Extracted RNA was then subjected to two rounds of DNase I treatment as follows to ensure degradation and removal of contaminating DNA. First, eluted RNA was added to a DNase Reaction Mix containing 40mM Tris-HCL (pH 7.9; Invitrogen), 6mM MgCl_2_ (Ambion), 10mM CaCl_2_ (Sigma) and 1 U DNase RQ1 (Promega) and incubated at 37°C for 15 minutes. Next, virion RNA was purified using the RNeasy Mini Kit with on-column DNase digestion with RNase-Free DNase I (Qiagen). The copies/μL in the virion standard were estimated by triplicate measurements using the Abbott RealTime SARS-CoV-2 assay (Abbott m2000 Molecular Platform). Dilutions of the virion standard were added to RT reactions to achieve expected inputs of 1 to 70,000 copies per 5uL RT (the input into each ddPCR well). RT reactions were performed as above, with random hexamers and poly-dT, except that the total volume of the RT was scaled up so that two replicate 5uL aliquots of cDNA could then be used to test each assay in parallel using replicate 20uL ddPCR reactions (see above) containing primers/probe specific for a given region.

### 2.5 Assay efficiency in presence of background RNA

Further validations were performed to determine each assay’s sensitivity to inhibition by “background” cellular RNA, as would be expected in clinical samples containing cells. The virion standard (1000 copies per 5μL RT) was added to RT reactions with or without cellular RNA from A549 cells (lung epithelial cell line) or donor PBMC (both added at 100ng/μl per RT, or 500ng per ddPCR well). RT reactions contained a total of 125μL with 12.5 μL of 10× SuperScript III buffer (Invitrogen), 12.5 μL of 50 mM MgCl2, 6.25 μL of random hexamers (50 ng/μL; Invitrogen), 6.25 μL of 50 μM dT15, 6.25 μL of 10 mM deoxynucleoside triphosphates (dNTPs), 3.125 μL of RNAseOUT (40 U/μL; Invitrogen), and 6.25 μL of SuperScript III RT (200 U/μL; Invitrogen). RT reactions were incubated at 25.0°C for 10 min, 50.0°C for 50 min, followed by an inactivation step at 85.0°C for 5 min. Undiluted cDNA (5 μL) was added to each 20 μL ddPCR reaction and replicate ddPCR reactions were performed for each assay.

### 2.6 Assay validations in clinical diagnostic samples from SARS-CoV-2 infected individuals

To investigate the viral transcription profile in clinical samples and determine whether our RT-ddPCR assays correlate with a clinical test, we obtained unused nucleic acid (ranging from 8.25-16.8μL) that remained after extraction by the Abbott m2000 instrument from nasopharyngeal swabs from 3 individuals who tested positive with the Abbott Real Time SARS-CoV-2 assay. Nucleic acid from these 3 individuals, who had C_t_ values of 11.59, 15.81, and 19.14 (respectively) on the Abbot assay, was tested using our RT-ddPCR assays for the 5’UTR, Main Proteinase, RdRp, S, M, N and 3’UTR regions. The available volume of nucleic acid was added into 85μL RT reactions containing 1× SuperScript III buffer, 5 mM MgCl2, 2.5 ng of random hexamers, 2.5 μM dT15, 0.5 mM deoxynucleoside triphosphates (dNTPs), 1U/μL of RNAseOUT, and 10U/μL of SuperScript III RT. RT reactions were performed under the aforementioned conditions. Undiluted cDNA was divided evenly across assays (5μL input into each ddPCR well, tested in duplicate) and ddPCR reactions were performed under the conditions described for ‘Validations using synthetic RNA’. Absolute values obtained by ddPCR were adjusted to account for differing input volume of nucleic acid to yield the SARS-CoV-2 copies/μL extract. The log-linear relationship between viral load measured by RT-PCR (Abbott Real Time SARS-CoV-2 assay) and RT-ddPCR was determined using GraphPad Prism (version 8.4.1).

## 3. Results

### 3.1 Detection limit, linearity, and efficiency using plasmid DNA

Two assays were designed for each region (indicated in Fig. 1; ‘Assay locations’) except the spike protein polybasic cleavage site. To evaluate the performance of each assay at the PCR stage, each pair of assays was tested on plasmid DNA. Since no commercially available plasmid contains the whole SARS-CoV-2 genome, and construction of such a plasmid is technically challenging (due to the 30kb length) and subject to higher biosafety restrictions, we constructed or purchased plasmids containing individual genes or regions. For each plasmid, the DNA concentration was measured by UV spectroscopy (NanoDrop) and the number of molecules (expected copies) was calculated using the molecular weight.

Each assay was assessed for detection limit, dynamic range, linearity, and efficiency by measuring the absolute number of copies detected using droplet digital PCR (ddPCR) from expected inputs of serially diluted plasmid DNA. All assays could detect as few as 1-10 copies and were linear over at least 3 orders of magnitude (R^2^>0.99 for all; Fig. 2). Assay efficiencies (measured by the slope) varied somewhat between assays, ranging from 0.67 (“N-ORF9_8”) to 1.1 (“M-ORF5”). One assay from each pair was selected for further study (Table 2; rejected primer/probe sets are listed in Table S1) based on the overall efficiency (Fig. 2), separation between the positive and negative droplets [amplitude/signal to noise] (Fig. S2), and specificity (Table S2). For the chosen assays, no positive droplets were detected with water or DNA from peripheral mononuclear blood cells (PBMC) from uninfected blood donors.

**Figure 2.**
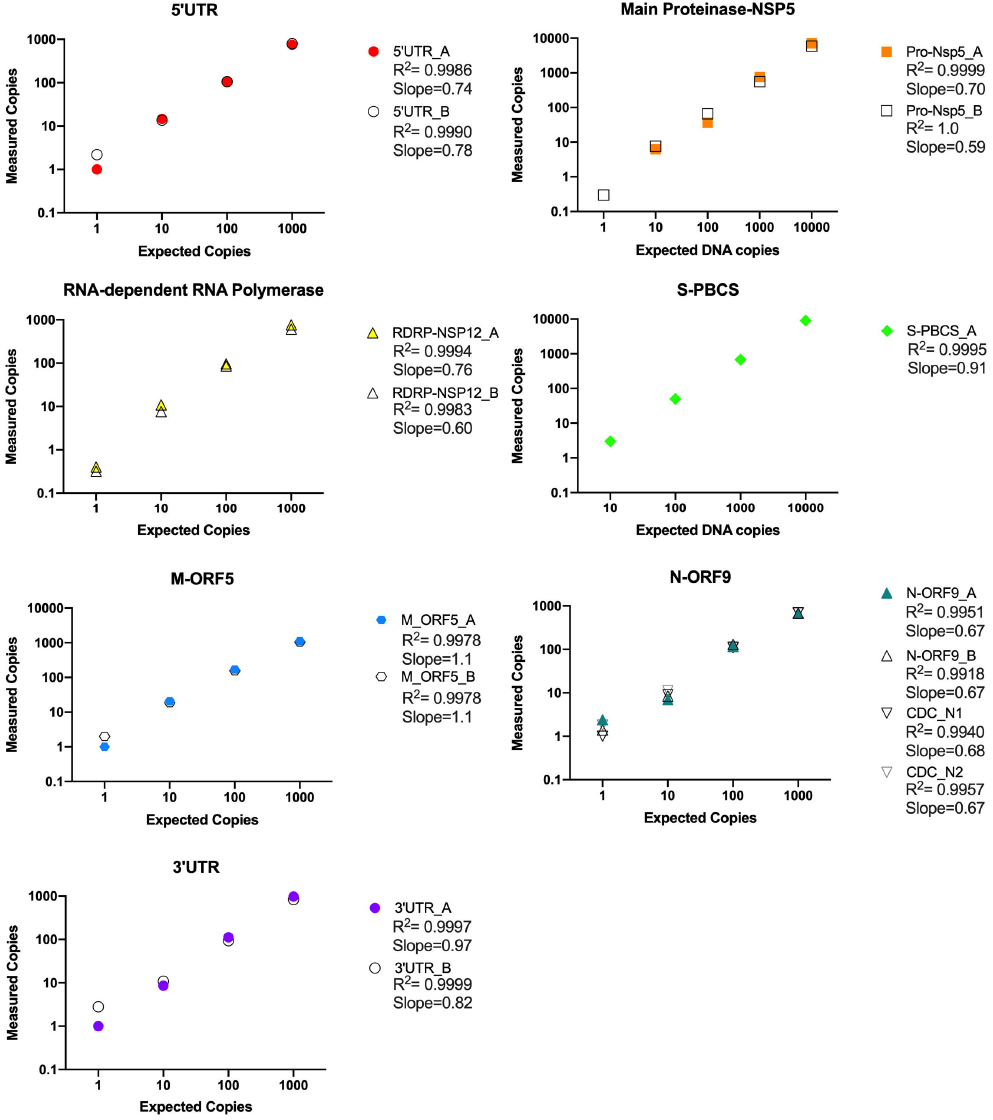
Efficiency and linearity of SARS-CoV-2 panel of ddPCR assays determined using plasmid DNA. Plasmids containing individual SARS-CoV-2 genes or regions were quantified by UV spectroscopy and diluted (expected copies) to test the absolute number of copies detected by each primer/probe set using duplicate ddPCR reactions (measured copies). Two primer/probe sets were tested for each region except the S-PBCS. One primer/probe set from each region (indicated by coloured symbol) was selected for subsequent experiments.

**Table 2.**
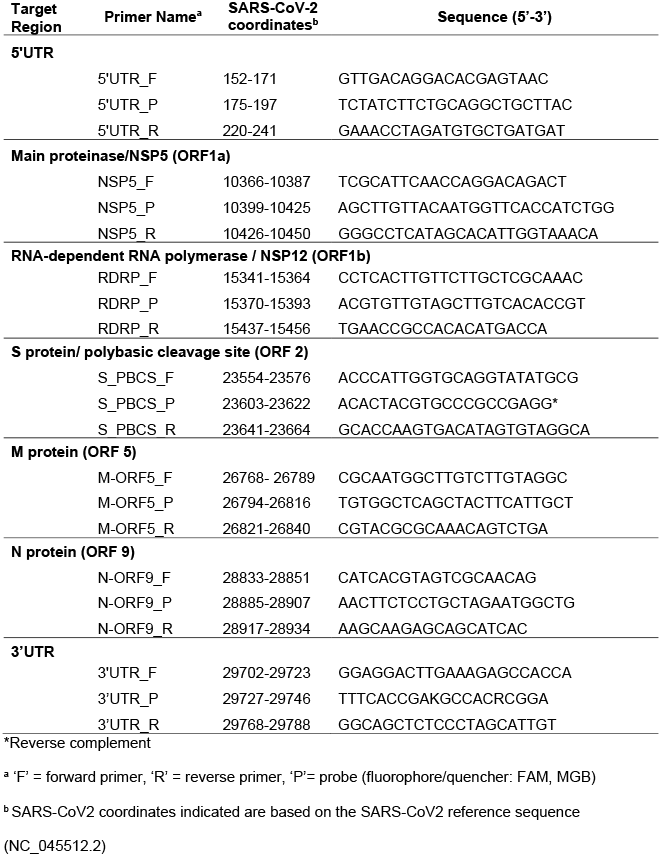
SARS-CoV-2 primer/probe sets selected for validation using IVT and virion RNA

### 3.2 Detection limit, linearity, and efficiency using in vitro transcribed and virion RNA

Selected assays for each region were tested using standards prepared from *in vitro* transcribed (IVT) RNA from the designed plasmids (5’UTR, Main Proteinase, RDRP, S, M, N and 3’UTR; Fig. 3 and Table 2). The expected copy numbers were calculated using the RNA concentration (as measured by UV spectroscopy [NanoDrop] and confirmed by the Agilent Bioanalyzer) and the molecular weight. Using RT-ddPCR, all assays could detect as few as 10 copies of RNA and demonstrated linearity over 3-4 orders of magnitude (R^2^>0.999 for all; Fig. 3). The efficiencies for detecting IVT RNA standards, which ranged from 0.18 (for Main Proteinase) to 0.96 (S-PBCS), were more variable than those observed for plasmid DNA. No amplification was detected in ‘No RT’ control reactions containing 10,000 IVT RNA copies/well, confirming the absence of any contaminating plasmid DNA. However, it is worth noting that none of these IVT standards were polyadenylated (so they should not be reverse-transcribed by poly-dT) and some of the standards were very short (<300 base pairs), which would likely limit the efficiency with which they were reverse transcribed by random hexamers. In addition, some of the measured differences in efficiency could reflect actual differences in the copy numbers present in the various IVT standards, which are difficult to determine precisely.

**Figure 3.**
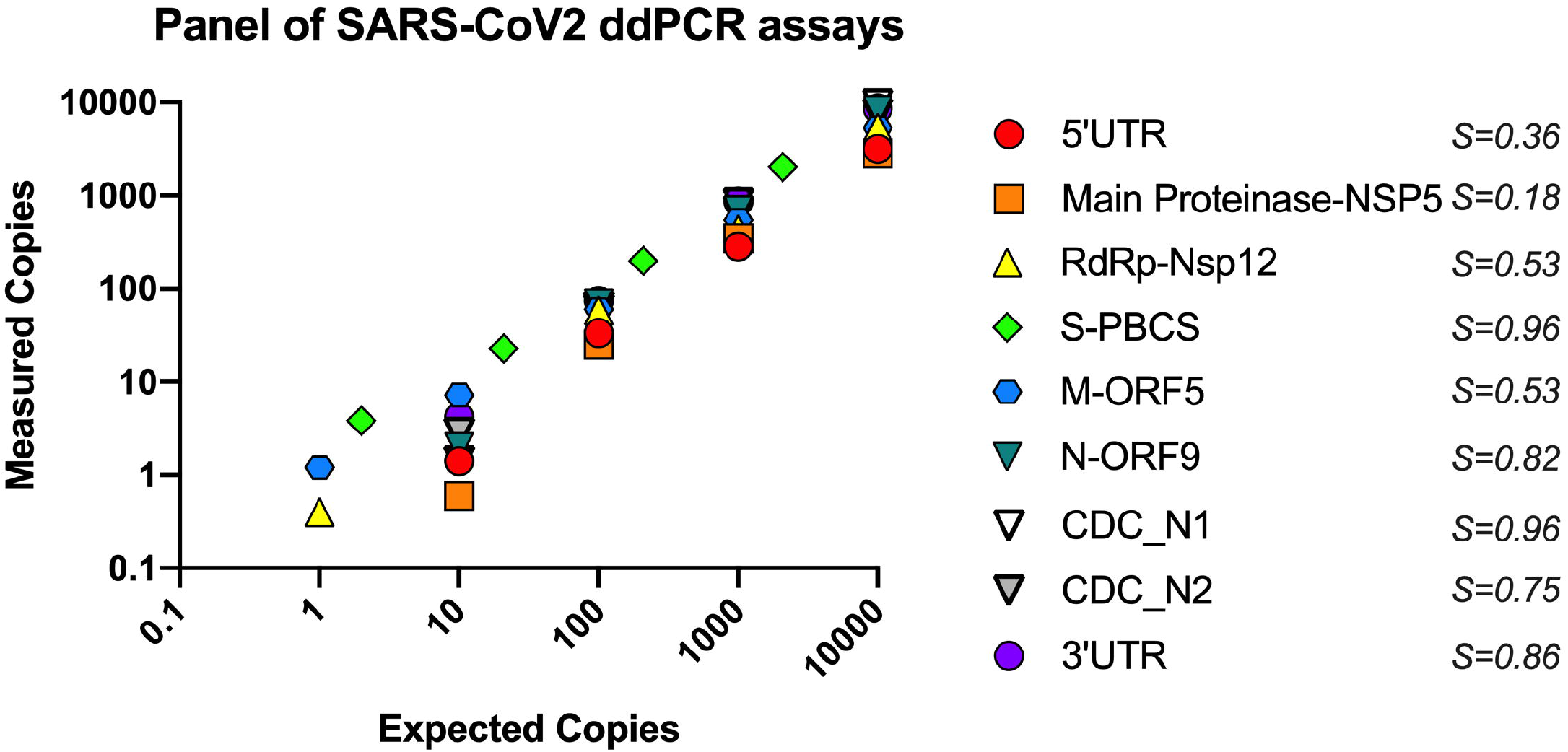
Efficiency and linearity of SARS-CoV-2 panel of ddPCR assays determined using in vitro transcribed (IVT) RNA. RNA standards containing a given region or gene of SARS-CoV-2 were prepared by in vitro transcription from plasmids and quantified by independent means (UV spectroscopy and the Agilent Bioanalyzer). Various inputs of each IVT RNA standard (which were used to calculate ‘Expected Copies’ per ddPCR well) were reverse transcribed and replicate aliquots of cDNA were used to measure the absolute number of copies detected by each ddPCR assay (‘Measured Copies’). Each assay was tested using expected inputs of 1-10^4^ copies per ddPCR well (except S-PBCS, which was tested at inputs of 2-2100 copies). Data represent average of duplicate wells from a representative experiment. *S*=slope, indicating assay efficiency. Each assay was tested in at least two independent experiments.

To circumvent these limitations, we prepared one SARS-CoV-2 ‘virion’ standard containing all of the target regions by extracting RNA from cell-free supernatant from a cell line (Vero CCL81) infected *in vitro* with a SARS-CoV-2 patient isolate (USA-WA1/2020). The expected copies in this virion standard were calculated using the C_t_ value measured by the Abbott m2000 Real Time SARS-CoV-2 viral load assay, which targets the N and RdRp genes using probes labelled with the same fluorophore. This virion standard enabled the preparation of common RT reactions containing specific inputs of SARS-CoV-2 genomic equivalents, from which aliquots of cDNA could be divided evenly across our panel of assays for simultaneous assessment of all target regions in ddPCR reactions (Fig. 4).

**Figure 4.**
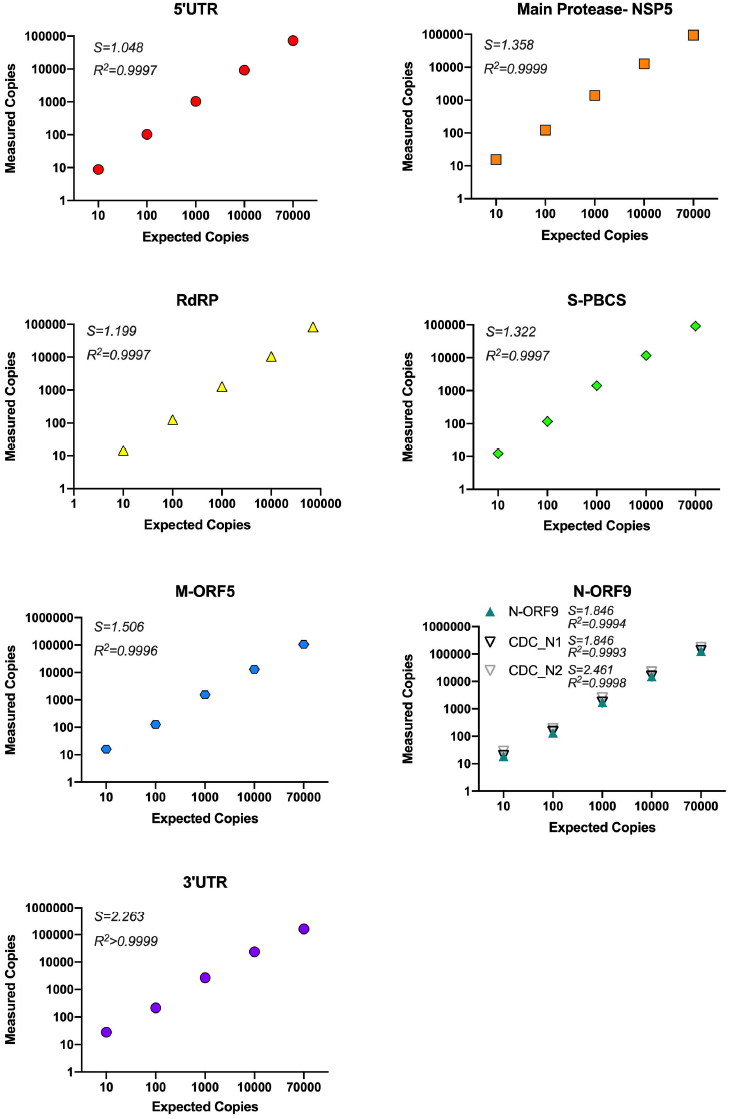
Efficiency and linearity of SARS-CoV2 panel of ddPCR assays determined using SARS-CoV-2 virion RNA. A SARS-CoV-2 “virion” standard was prepared by extracting the RNA from the supernatant of an in vitro infection and quantified using the Abbott Real Time SARS-CoV-2 assay. Various inputs of the virion standard (which were used to calculate ‘Expected Copies’ per ddPCR well) were applied to a common reverse transcription reaction, from which aliquots of cDNA were used to measure the absolute number of copies detected by each ddPCR assay (measured copies). Each assay was tested with expected inputs of 10-10^4^ copies/ddPCR well in duplicate. S (slope) and R^2^ are indicated for each assay. Representative data for n=2 independent experiments are shown.

Expected inputs of 10 to 70,000 copies per well were used to measure the absolute copies of 5’UTR, Main Proteinase, RdRp, S, M, N and 3’UTR regions. All assays detected as few as 10 copies of the virion standard and were linear over four orders of magnitude (R^2^>0.999 for all). No amplification was detected in ‘No RT’ control reactions containing 10,000 IVT RNA copies/well. Assay efficiencies were all greater than 1.0 (range: 1.05 to 2.46), likely because the estimate from the Abbott assay was lower than the true value and/or the RT-ddPCR assays are more efficient. In addition, the efficiency of the RT-ddPCR assays increased from 5’ to 3’ targets, which could reflect the presence of 3’ subgenomic RNAs in the virion standard or greater efficiency of reverse transcription from the 3’ end of the genome.

### 3.3 Assay specificity and false positive rate

To determine the non-specific reactivity of oligonucleotides (false positive rate) for each assay, we performed a median of 26 [range 18-32] ‘no template’ controls (NTC). These reactions were performed with both water (water NTC) and DNA or RNA isolated from SARS-CoV-2-negative donor PBMC (DNA/RNA NTC) (Table S2). Except for one experiment using IVT RNA, where a total of three droplets were detected across duplicate NTC wells containing donor PBMC tested for Main Proteinase-NSP5, no other false positives were observed.

### 3.4 Comparison of new and existing SARS-CoV-2 assays in ddPCR platform

Our assay panel included new primers/probes for the nucleocapsid (N-ORF9), which is targeted by existing diagnostic real-time PCR assays. We compared the performance of our ‘N-ORF9’ primers/probe to the primers/probes from the U.S. Center for Disease Control assays for the nucleocapsid (CDC-N1 and CDC-N2)^31^ using ddPCR. The N-ORF9 assay efficiency was similar to that of CDC-N1 and CDC-N2 for plasmid DNA, in between that of CDC-N1 and CDC-N2 for IVT RNA, and similar to CDC-N1 for the virion standard (Fig. 2-4).

In addition, we compared our primers/probes for the RdRp to published primers/probes for the “IP2” assay^32^ (which targets ORF1a) and “E-Sarbeco”^33^ assay (which targets the E gene) using RT-ddPCR and the virion standard (Fig. 5; Table 3). The IP2 (ORF1a) assay efficiency was 1.11, compared to 1.20-1.28 for our RdRp (ORF1b) and 1.36 for our main protease (ORF1a) assays (Fig. 4-5). The E-Sarbeco [ORF4] assay efficiency (1.08) was similar to the IP2, but may have been less than our assays targeting neighboring genes (S-PBCS [ORF2]: 1.32; M-ORF5: 1.51).

**Figure 5.**
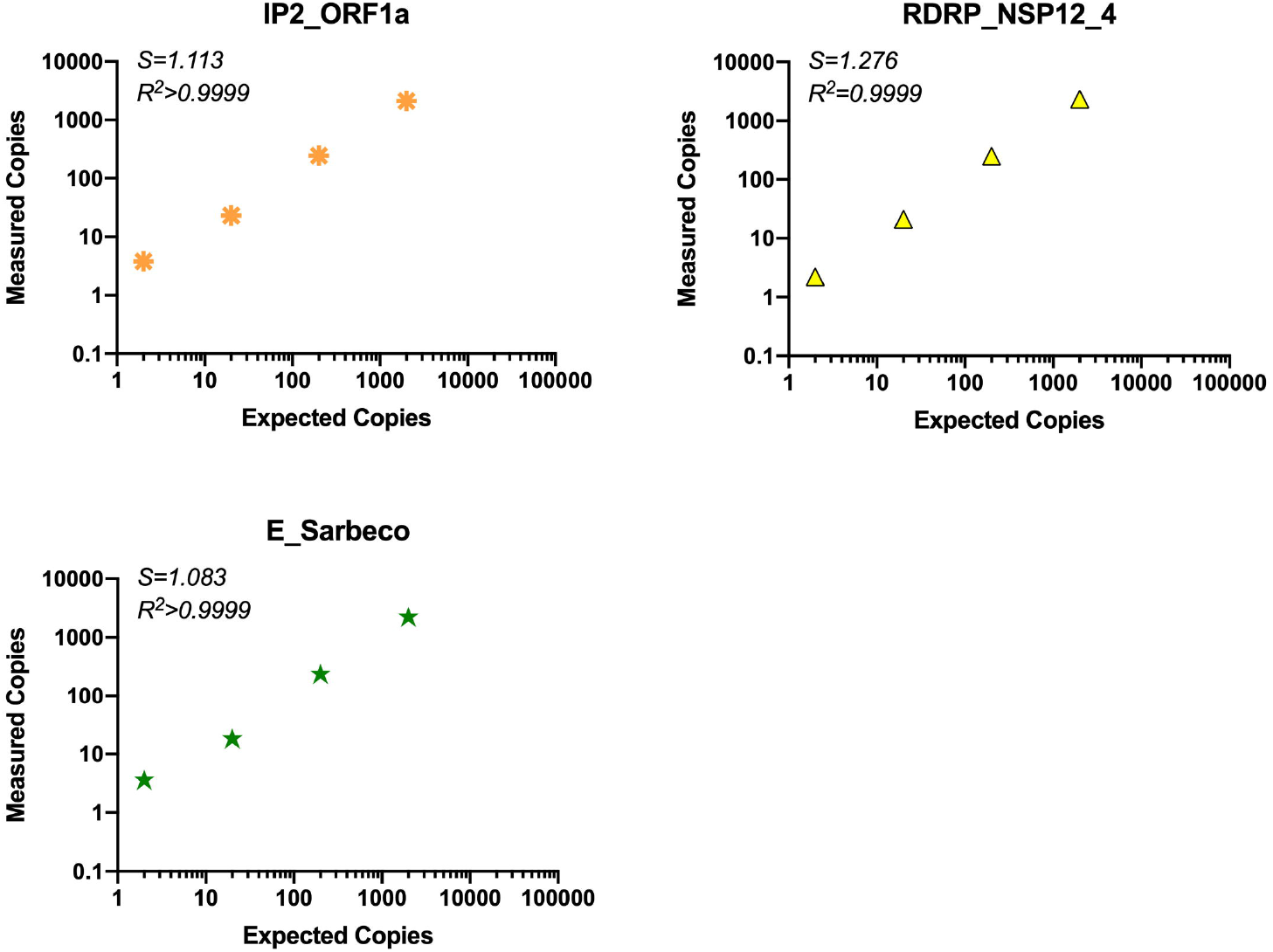
Comparison of assay efficiency and linearity of published assays, ORF1a “nCoV_IP2” and E gene and novel RDRP-NSP12 assay. The performance of our RDRP-NSP12 assay was compared to published primers/probes for ORF1a and the E gene in the ddPCR platform using common RT reactions containing virion standard RNA inputs of 2-2×10^4^ copies/ddPCR well. S (slope) and R^2^ are indicated for each assay.

**Table 3.**
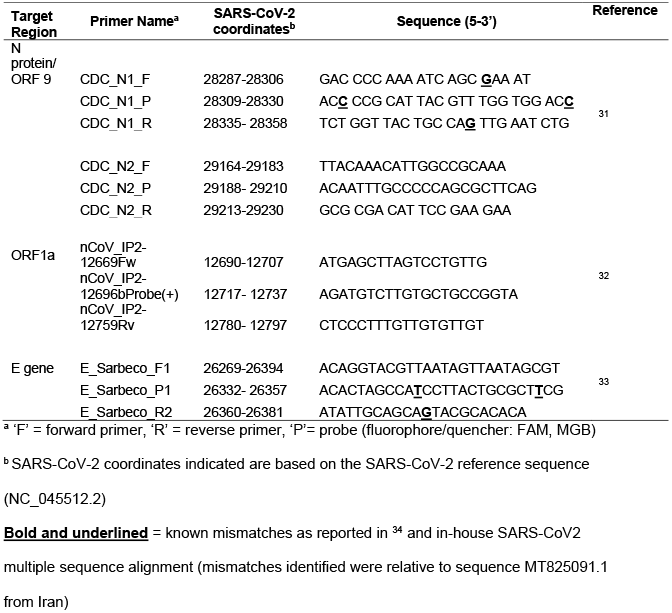
SARS-CoV-2 assays from other sources

### 3.5 Lower limit of detection of SARS-CoV-2 in RNA

Our validation studies included SARS-CoV-2 RNA inputs down to 1 copy per ddPCR reaction (Fig. 2-3). We estimated the lower limit of detection (LLOD) for each assay in our panel based on data for all replicates tested at 10 copy and 1 copy inputs (Table S3). At 10 copies, all of our assays detected SARS-CoV-2 in ≥85.7% of tests (range= 85.7-100%). At 1 copy input, our assays detected SARS-CoV-2 in ≥25% of tests (range=25-88%), underscoring the high sensitivity of our assays.

### 3.6 Effect of Background RNA on assay efficiencies

Next, we assessed the efficiencies of our assays in the presence of “background” RNA from uninfected cells (Fig. 6). At a constant input of 1000 copies of the SARS-CoV-2 virion RNA, we determined the effect of adding cellular RNA (100ng per μL of RT) extracted from PBMC or a lung epithelial cell line (A549 cells). All assays showed slightly greater efficiency in the presence of 100ng/μL background RNA from either PBMC or A549 cells compared to the virion standard with no background RNA. No false positives were detected with 100ng/μL RT RNA from PBMC, while 1-4 droplets were sometimes detected in the RNA from A549 cells using some assays (Main Proteinase, RdRp, S-PBCS). Overall, these data suggest that in samples derived from individuals infected with SARS-CoV-2, our assays are likely to be minimally inhibited by background RNA, making them ideally suited to a diverse range of clinical samples.

**Figure 6.**
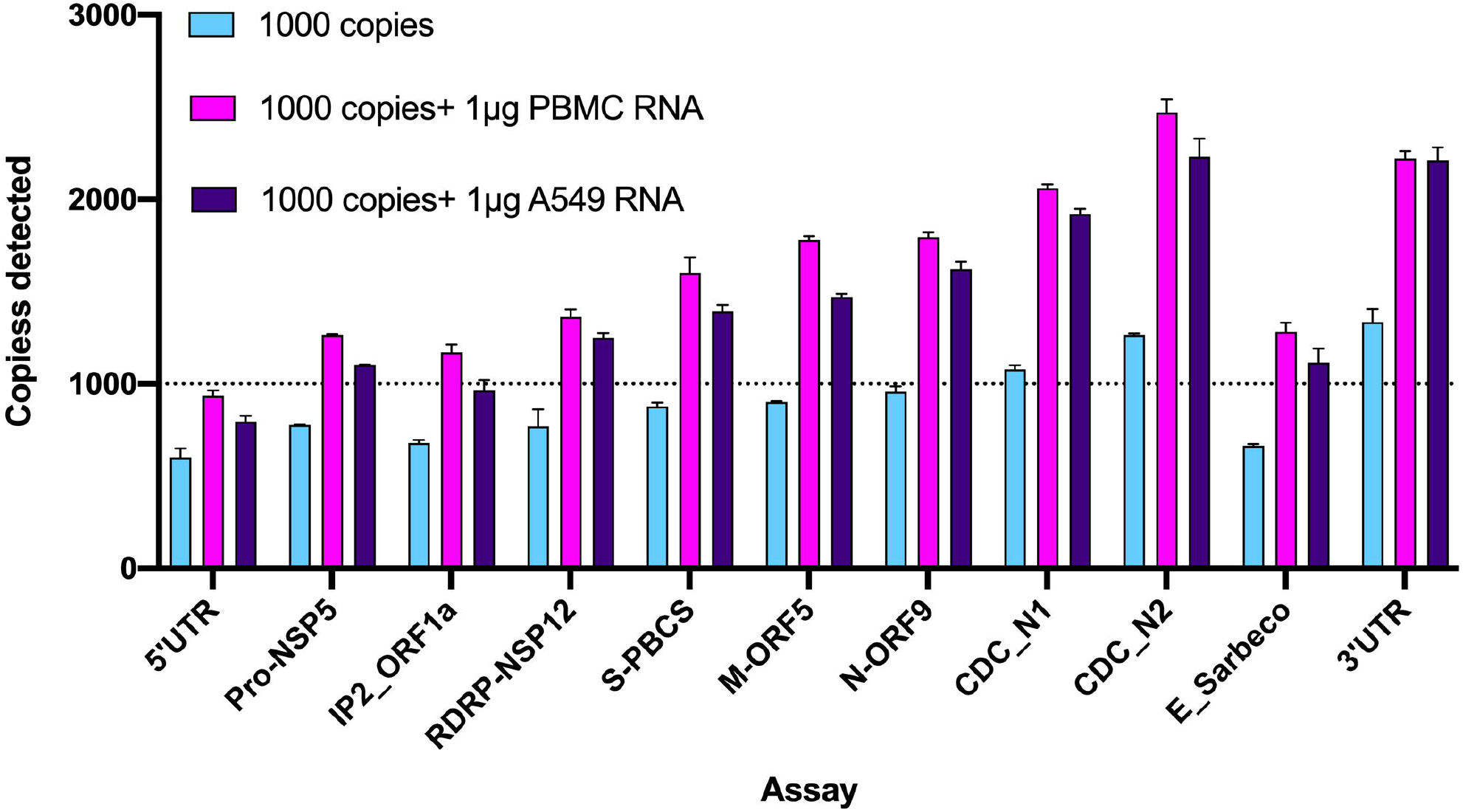
Effect of background RNA on ddPCR assay performance. We simultaneously tested all assays in our panel against reported assays, CDC_N1, CDC_N2, E_Sarbeco, and IP2_ORF1a, in the presence and absence of background RNA. Each assay was tested with a constant input of SARS-CoV-2 virion standard (predicted to yield 1000 copies/ddPCR well) in the presence or absence of background RNA from PBMC or a lung epithelial cell line (A549) added at a concentration of 100ng/μL of RT reaction (500 ng/ddPCR well, or 1 μg for the 2 replicate wells used to test each assay). Negative controls included water, 1 μg/assay PBMC RNA, and 1 μg/assay A549 RNA. Assays are indicated on x-axis in order from 5’ to 3’ and dotted line indicates 1000 SARS-CoV-2 RNA copy input. Error bars represent standard deviation from duplicate wells.

### 3.7 Strong correlation between viral loads measured by RT-ddPCR and real-time PCR in clinical diagnostic samples

To compare our assays with a clinical test, we obtained unused nucleic acid that had been extracted by the Abbott m2000 molecular platform from nasopharyngeal swabs from three SARS-CoV-2-infected individuals and remained after clinical testing using the Abbott Real Time SARS-CoV-2 assay. Using this nucleic acid, we measured RNA levels of the 5’UTR, Main Proteinase, RdRp, S, M, N and 3’UTR regions using our RT-ddPCR assays. (Fig. 7). As observed with the virion standard, transcripts containing the most 3’ regions (N-ORF9 and 3’LTR) tended to be present at higher copy numbers, while those containing the 5’LTR tended to be present at lower levels. However, the order of transcript levels varied somewhat between individuals and sometimes differed from the 3’ to 5’ gradient observed with the virion standard. For example, levels of S-PBCS RNA tended to be lower than those of the more 5’ Main Protease (NSP5) transcripts. These potential differences in SARS-CoV-2 transcription profile may reflect changes in viral dynamics over the course of infection or inter-individual variability in viral sequences or host responses, and should be confirmed in future studies using longitudinal samples from more individuals.

**Figure 7.**
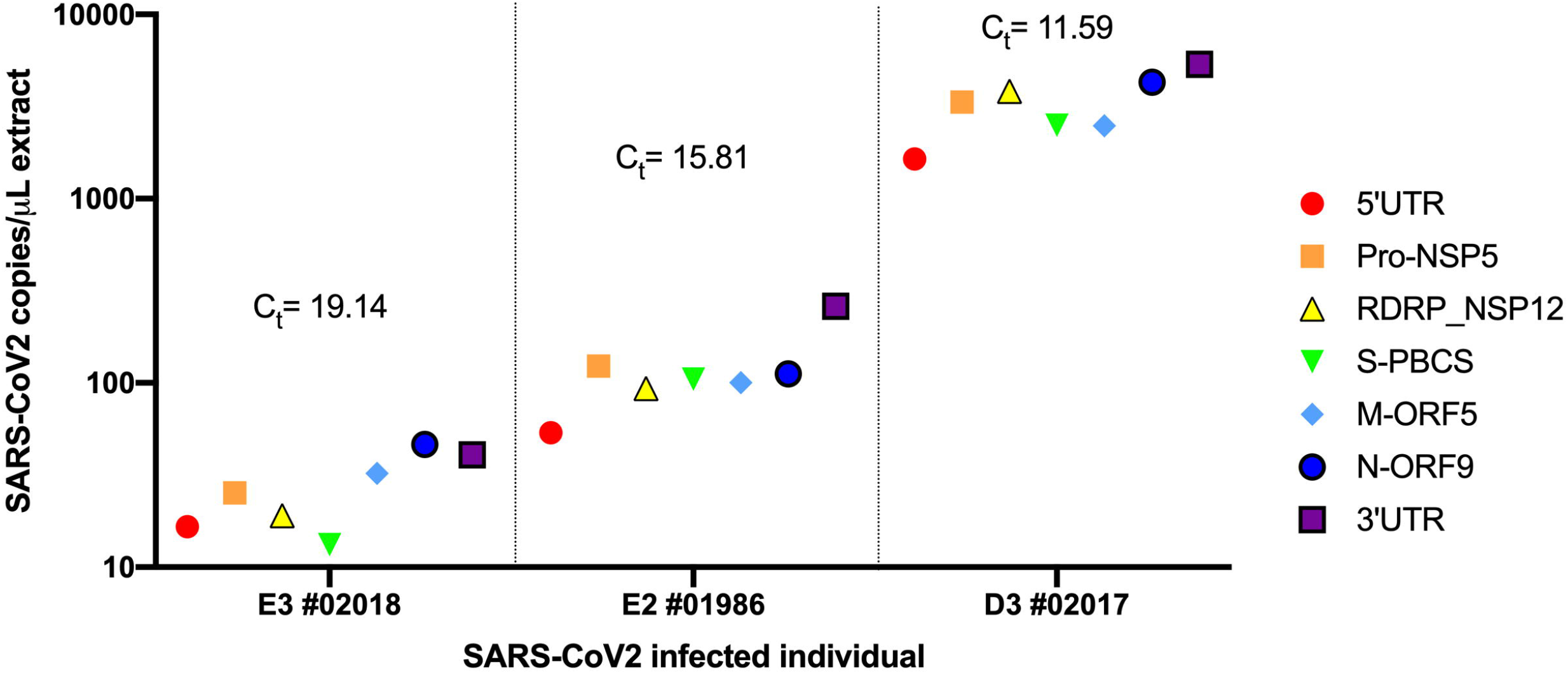
Transcription profile of three SARS-CoV-2 infected individuals determined using RT-ddPCR. Unused nucleic acid (ranging from 8.6-16.8 μL) extracted by the Abbott m2000 platform from nasopharyngeal swabs from SARS-CoV-2 infected individuals (n=3) was used in a common RT reaction for each individual. Resulting cDNA was divided evenly across reactions for the seven assays in our panel and targets were measured using ddPCR. Colored symbols indicate SARS-CoV-2 target region. Copy numbers from each assay are expressed as SARS-CoV-2 copies per μL of nucleic acid and grouped for each individual (x-axis). Threshold cycle (C_t_) values, as determined by Abbott Real Time SARS-CoV-2 viral load assay, are indicated above each individual’s dataset.

Next, we determined the correlation between the C_t_ value as measured by the Abbott assay and SARS-CoV-2 copy numbers as determined by RT-ddPCR. For each target, this relationship was modelled using linear regression following log transformation of SARS-CoV-2 copies/μL extract [where y=Log_2_(x)] (Fig S1). The coefficient of determination (R^2^) for each model was ≥0.93 for all targets, underscoring the log-linear relationship between ddPCR-based SARS-CoV-2 transcript levels and C_t_ values in diagnostic specimens. Taken together, these data strongly underscore the sensitivity of our assays, demonstrating the ability to detect all targets using minimal RNA inputs (effectively 1.2-2.4 μL RNA input per assay), and their strong correlation with C_t_ values obtained by real-time PCR using clinical assays. Furthermore, these data highlight that delineation of the SARS-CoV-2 transcription profile in samples across differing timepoints within and between participants may yield valuable insight into viral transcription dynamics across the course of SARS-CoV-2 infection.

## 4. Discussion

The 2019 SARS-CoV-2 outbreak has heralded the development of an array of diagnostic molecular tools to study this novel coronavirus. However, currently described PCR-based diagnostic assays are qualitative or semi-quantitative, are limited to the simultaneous detection of one or two regions, and do not distinguish genomic from subgenomic RNAs. Here, we report a panel of new primer/probe sets that span the SARS-CoV-2 genome and target important nongenic regions, non-structural genes found only in genomic RNA, and structural genes that are also found in different subgenomic RNAs.

We used these new primers/probes for RT-ddPCR rather than qRT-PCR because ddPCR provides absolute quantification (does not require an external calibrator), tends to tolerate sequence mismatches in primer/probe sequences better than qRT-PCR, and may be more precise at low copies, while providing similar sensitivity and reproducibility^30,35^. During validation of these assays with multiple different standards, we sometimes found that the efficiency of the same assay varied somewhat across different standards. These differences may reflect differences in the nature of the standards (DNA, short *in vitro* transcribed nonpolyadenylated RNA, or “virion RNA”) as well as the difficulty in determining the exact number of copies in an external standard; the latter issue highlights a major advantage of the absolute quantification provided by ddPCR. On all standards tested, the seven RT-ddPCR assays were extremely sensitive (down to 1-10 copies) and linear over 3-4 orders of magnitude, and all seven assays showed no inhibition by up to 500,000 cell equivalents of RNA per ddPCR well, suggesting that these assays could be extremely useful for SARS-CoV-2 research. While most existing clinical assays for SARS-CoV-2 use qPCR because it is less expensive and may have fewer false positives than ddPCR, it is likely that the primer/probe sets described here would also work well in qPCR assays for research or clinical testing.

The utility of assays that target multiple genomic regions is supported by studies demonstrating loss in sensitivity of published assays owing to mutations that could affect primer annealing. For instance, a recent study found that 34.38% (11,627) of SARS-CoV-2 genomes featured a single mutation capable of affecting annealing of a PCR primer in tested assays from the World Health Organization, Centers for Disease Control and Prevention, National MicrobiologyData Center, and Hong Kong University^36^. Another study found single nucleotide mismatches in 0.2% and 0.4% of the surveyed SARS-CoV-2 sequences compared to the CDC-N1 probe and reverse primer, respectively, and 0.4% of those sequences compared to Charité’s E_Sarbeco_R primer^34^. Therefore, a strategy that can target multiple genomic regions may have utility in sensitive detection of SARS-CoV-2.

Extensive, well-designed studies have assessed the analytical sensitivity and efficiency of existing RT-qPCR primer-probes sets^34,37–39^ and explored adaptation of such assays to the ddPCR platform^40^. In this study, we describe how some of the available diagnostic assays compare to our novel SARS-CoV-2 assays and report how a multi-assay approach using the ddPCR platform could significantly advance our understanding of SARS-CoV-2 transcription and replication. While highly-sensitive PCR-based assays might not be essential to identify SARS-CoV-2-infected individuals in the transmissible/contagious phase of infection, quantitative assays capable of detecting very low copies of SARS-CoV-2 will be particularly useful in understanding the course of infection and correlates of disease progression. Existing clinical assays are quite sensitive for detecting COVID-19 during the first several weeks of infection, but often become negative after 2-3 weeks of infection^41–43^. In some cases, individuals who test positive may have a subsequent negative test followed by another positive or alternating positive and negative tests^44,45^. Some individuals may also have prolonged viral shedding after symptomatic relief, with one study noting a patient with qRT-PCR positivity detected in upper respiratory tract samples 83 days post-symptom onset^46^. Therefore, sensitive assays such as those described in the study could be of great utility in studying the course of infection two or more weeks after the resolution of acute symptoms. Another advantage of the approach described here is that it permits a single sample to be simultaneously assayed for multiple targets, which may increase sensitivity and specificity while helping to delineate the transcriptional profile of SARS-CoV-2 in infected patient samples. As such, this panel of assays can be applied to a diverse range of clinically relevant samples in which SARS-CoV-2 RNA may be in low or high abundance.

Using both the virion standard and clinical samples from the nasopharynx, we tended to observe higher copy numbers for targets at the 3’ end of the genome (N, 3’UTR) compared to the 5’ end (5’UTR, main protease). This discrepancy is not explained by differences in PCR efficiency, since the efficiency of the N assay on plasmid DNA was actually lower than that of assays for the 5’UTR or main protease. It is possible that reverse transcription is more efficient for assays at the 3’ end (perhaps due to more efficient reverse transcription from the poly-dT), although random hexamers should bias towards the 5’ end and the combination has been used to prevent bias towards either the 5’ or 3’ end of the 9.6kb genome of HIV-1^30,47^. It is also possible that the 3’ assays measure higher copies because they are detecting subgenomic RNAs generated by infected cells and not packaged into virions^48^, which may have been present in the supernatant used to prepare the virion standard if they were released from dying cells or present in low levels in exosomes. This excess of targets corresponding to sgRNAs may be much greater in samples that contain more cells or cell-associated RNA, and it has important implications for clinical testing and research. For targets in the 3’ third of the genome that are transcribed as sgRNAs, regions that are further downstream (3’) may be incorporated into a greater variety of sgRNAs and therefore should be present at higher copy numbers, so assays targeting these regions may be more sensitive to detect infection^8^. On the other hand, sgRNAs are not infectious, so assays targeting more 5’ regions that are transcribed only as genomic RNA (ORF 1a and 1b) may correlate better with infectivity.

The clinical implications of SARS-CoV-2 subgenomic RNA transcription are currently unknown. The synthesis of subgenomic RNAs is a common strategy employed by positive-sense RNA viruses to transcribe their 3’ proximal genes that encode products essential for particle formation and pathogenesis^49–51^. In coronaviruses such as mouse hepatitis virus (MHV), the synthesis of subgenomic RNAs may function as important mediators of positive strand synthesis^52^, and more broadly, members of the order *Nidovirales* (*including Coronaviridae*) feature high levels of redundancy to ensure continued protein synthesis even in the event of point mutations in regulatory sequences^53^. The characterization of the SARS-CoV-2 transcription profile in differing patient samples over the course of infection may provide insight into the molecular mechanisms by which SARS-CoV-2 regulates gene expression through differential transcription of genomic and subgenomic RNAs, and how this differential gene expression may contribute to pathogenesis.

We found that our assays performed better in the presence of background RNA, irrespective of origin (blood or epithelial cells, Fig. 6). This finding accords with other studies that have extensively validated the effect of differing variables on RT efficiency and suggest that the presence of some background RNA may increase efficiency of the reverse transcription step^54–56^. While the efficiency of our assays tended to decrease with RNA concentrations above 100ng/μL RT, even at 500ng RNA/μL RT, these assays still performed better than in the absence of any background RNA, suggesting that they are ideally suited for testing samples from different tissues where the levels of genomic RNA may differ considerably. Furthermore, our comparison of viral loads obtained by RT-ddPCR and qRT-PCR demonstrates the strong correlation between data obtained from these two platforms and the minimal RNA input required to yield robust data using our RT-ddPCR assays.

Limitations of this study should be acknowledged. In order to test our assays in parallel with published assays (total of 11 assays) in background RNA experiments (Fig. 7), we increased RT reaction volumes from 50-70 μL to 125 μL to accommodate the additional assays. In the absence of background RNA, the efficiency appeared to be higher in the 50-70μL RT reactions (Fig. 4-5, >100% efficiency for all assays) than the 125μL reactions (Fig. 7; median efficiency=88% [range: 60-133%]). If the discrepancy is not due to a difference in the actual input of the standard, it is possible that larger reaction volumes lead to less efficiency in reverse transcription. However, for application to patient samples, our core panel of 7 assays (Table 1) is sufficient to provide a detailed view of the transcription profile of SARS-CoV-2, so preparation of RT reactions >70μL will likely be unnecessary.

For our study of the viral transcription profile and correlation with the C_t_ value as determined by the Abbott SARS-CoV-2 Real Time Assay, a limited amount of nucleic acid was available from only a small number of de-identified individuals. Despite this small sample size, we demonstrated both the sensitivity of all assays in our panel and their strong correlation with C_t_ values in diagnostic specimens. These data allude to potential differences in the transcription dynamics of SAR-CoV-2 during the course of infection and merit further investigation.

## Conclusions

We developed a panel of sensitive, quantitative RT-ddPCR-based SARS-CoV-2 assays that collectively span the genome and target nongenic and genic regions, genes encoding for important enzymes and structural proteins, and genes found in different subgenomic RNAs. These assays can serve as novel molecular tools to investigate SARS-CoV-2 infection, replication dynamics, and gene expression to better understand the viral dynamics and pathogenesis of SARS-CoV-2 over the course of infection. Future studies employing these assays will enhance our understanding of SARS-CoV-2 replication and transcription and may also inform the development of improved diagnostic tools and therapeutics.

## Additional Information

### Ethics

This study included the use of de-identified nucleic acid from three SARS-CoV-2-infected individuals. The study authors had no subject contact or access to any personally-identifiable information (Category 4, IRB exempt).

### CRediT authorship contribution statement

**Sushama Telwatte**: Conceptualization, Data curation, Formal analysis, Funding acquisition, Investigation, Methodology, Supervision, Validation, Visualization, Writing - original draft. **Nitasha Kumar**: Investigation, Writing - review & editing. **Chuanyi M. Lu**: Resources, Writing - review & editing. **Alberto Vallejo-Gracia**: Investigation, Writing - review & editing. **G. Renuka Kumar**: Investigation, Writing - review & editing. **Melanie Ott**: Resources, Supervision, Writing - review & editing. **Joseph K. Wong**: Resources, Supervision, Writing - review & editing. **Steven A. Yukl**: Conceptualization, Funding acquisition, Methodology, Resources, Supervision, Writing - original draft.

