## Supplementary Figures and Tables for "Novel RT-ddPCR Assays for determining the transcriptional profile of SARS-CoV-2"

**Table S1. Primer/probe sets that were not selected following validations with plasmid DNA**

| Target Region | Primer Name[a] | SARS-CoV-2 coordinates[b] | Sequence (5'-3') |
|---|---|---|---|
| **5'UTR** | | | |
| | 5'UTR_B_F | 185-259 | GCAGGCTGCTTACGGTTTCG |
| | 5'UTR_B_P | | CCGTGTTGCAGCCGATCATCAGC |
| | 5'UTR_B_R | | TTTCGGTCACACCCGGACGA |
| **Main proteinase/NSP5 (ORF1a)** | | | |
| | NSP5_B_F | 10612-10689 | TGACAGGCAAACAGCACAAGC |
| | NSP5_B_P | | AGCTGGTACGGACACAACTATTACAGT |
| | NSP5_B_R | | ACAGCAGCGTACAACCAAGC |
| **RNA-dependent RNA polymerase / NSP12 (ORF1b)** | | | |
| | RDRP_B_F | 15437-15551 | TGGTCATGTGTGGCGGTTCA |
| | RDRP_B_P | | ACCAGGTGGAACCTCATCAGGA |
| | RDRP_B_R | | ACATTGGCCGTGACAGCTTG |
| **M protein (ORF 5)** | | | |
| | M-ORF5_B_F | 26885-26965 | CGTGCCACTCCATGGCACTAT |
| | M-ORF5_B_P | | TGACCAGACCGCTTCTAGAAAGTGA |
| | M-ORF5_B_R | | TGTCCACGAAGGATCACAGCTC |
| **N protein (ORF 9)** | | | |
| | N-ORF9_B_F | 28471-28632 | CCCTCGAGGACAAGGCGTTC |
| | N-ORF9_B_P | | ACCGTCACCACCACGAATTCGT* |
| | N-ORF9_B_R | | CCAGCTTCTGGCCCAGTTCC |
| **3'UTR** | | | |
| | 3'UTR_B_F | 29708- 29805 | CTTGAAAGAGCCACCACAT |
| | 3'UTR_B_P | | ACAGTGAACAATGCTAGGGAGAG |
| | 3'UTR_B_R | | TAGGGCTCTTCCATATAGGC |

**Table S2. False positive data from no template controls**

| No template control | Assay[a] | | | | | | |
|---|---|---|---|---|---|---|---|
| | **5'UTR** | **Main Proteinase-NSP5** | **RdRP** | **S-PBCS** | **M-ORF5** | **N-ORF9** | **3'UTR** |
| **$H_2O$** | 0/16 | 0/12 | 0/13 | 0/9 | 0/10 | 0/14 | 0/14 |
| **PBMC DNA** | 0/2 | 0/2 | 0/2 | 0/2 | 0/2 | 0/2 | 0/2 |
| **PBMC RNA** | 0/16 | 2/12* | 0/13 | 0/9 | 0/10 | 0/14 | 0/14 |

[a]For each assay, number of wells in which at least one droplet was detected is shown.

*A total of 3 droplets were detected in two wells from the same assay.

**Table S3. Frequency of detection of SARS-CoV2 for each ddPCR assay**

| Input (Copies) | % replicates in which signal was detected | | | | | | | | |
|---|---|---|---|---|---|---|---|---|---|
| | 5'UTR | Main Pro-NSP5 | RDRP | S | M-ORF5 | N-ORF9_8 | CDC_N1 | CDC_N2 | 3'UTR |
| 10 | 92.9 | 85.7 | 88.9 | 100.0 | 88.9 | 92.9 | 92.9 | 100.0 | 100.0 |
| 1 | 43.8 | 30.0 | 25.0 | 87.5 | 25.0 | 37.5 | 25.0 | 30.8 | 31.3 |

Each assay was tested at the given copy input in an average of 14 replicates/assay.

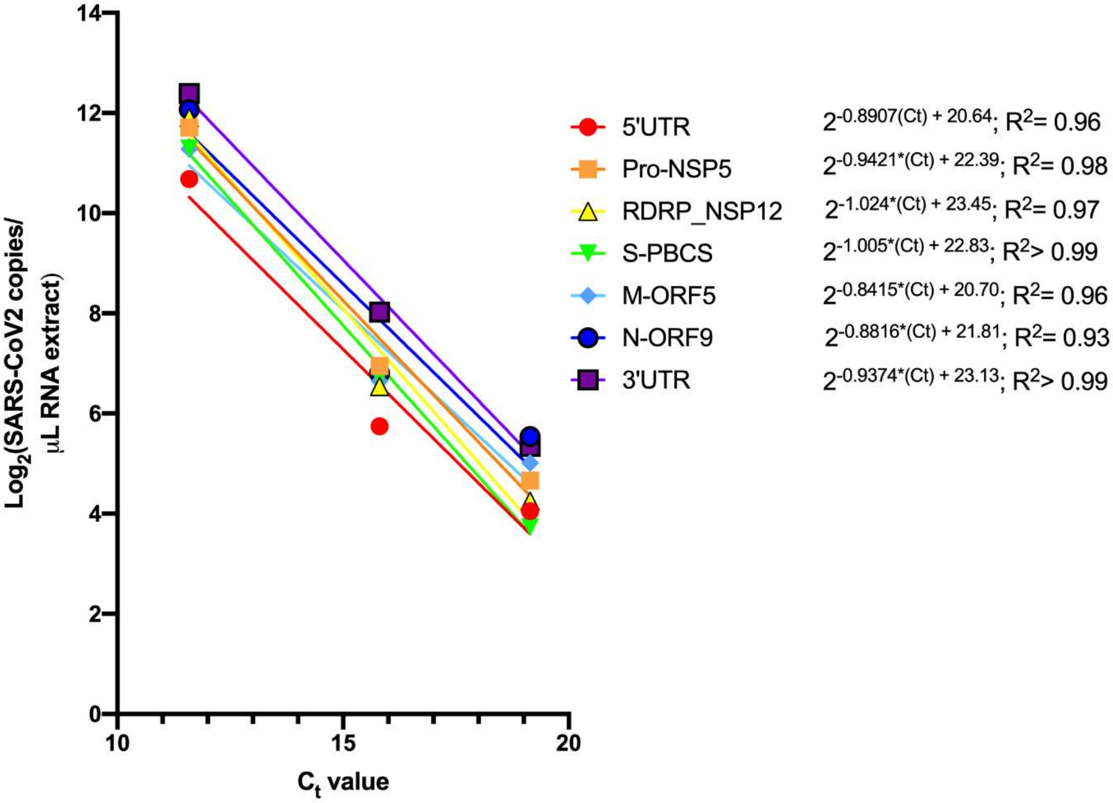


**Fig S1. Relationship between $C_t$ and viral load as determined by RT-ddPCR.**

Expression of seven SARS-CoV-2 targets [log2(SARS-CoV-2 copies/µL RNA extract)] was plotted relative to Ct value determined by Abbott SARS-CoV-2 Real Time viral load assay. We observed a strong correlation between these two variables. We modelled the relationship between the $C_t$ value and ddPCR-based viral load using linear regression (GraphPad Prism; version 8.4.1) and determined equations that describe the log-linear relationship between these two variables for each target in our panel.

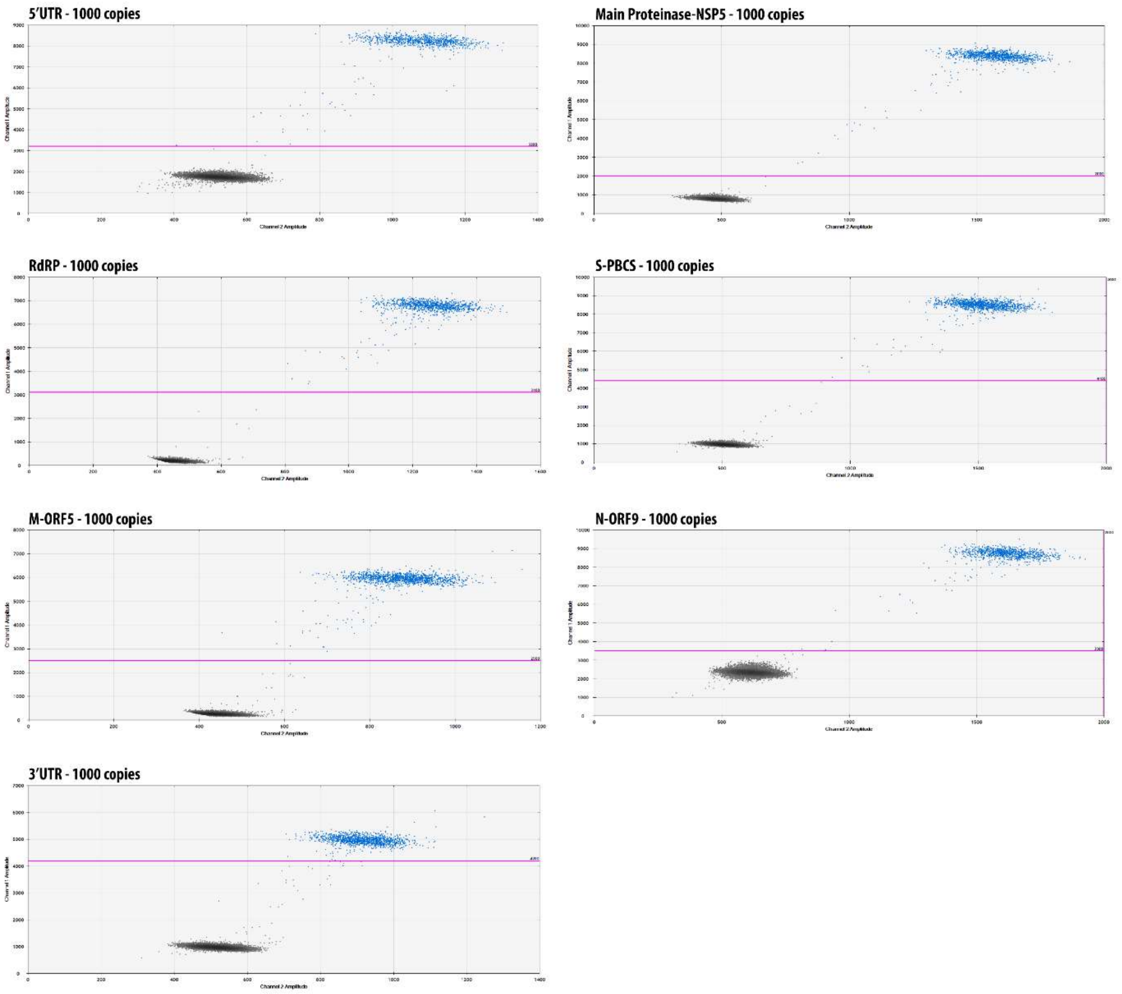


**Fig S2. Two dimensional ddPCR plots for SARS-CoV-2 assay panel.** Plasmid DNA at 1000 copies per ddPCR well are shown. Droplets were read and analyzed using the QuantaSoft™ software in the 'absolute quantification' mode. Primer/probes sets comprising our final panel, for which the difference in amplitudes of positive and negative droplets were higher than other primer/probe sets tested, are shown. Plots are representative of signal to background observed for each assay.
